# FlyLimbTracker: an active contour based approach for leg segment tracking in unmarked, freely behaving *Drosophila*

**DOI:** 10.1101/089714

**Authors:** Virginie Uhlmann, Pavan Ramdya, Ricard Delgado-Gonzalo, Richard Benton, Michael Unser

## Abstract

Understanding the biological underpinnings of movement and action requires the development of tools for precise, quantitative, and high-throughput measurements of animal behavior. *Drosophila melanogaster* provides an ideal model for developing such tools: the fly has unparalleled genetic accessibility and depends on a relatively compact nervous system to generate sophisticated limbed behaviors including walking, reaching, grooming, courtship, and boxing. Here we describe a method that uses active contours to semi-automatically track body and leg segments from video image sequences of unmarked, freely behaving *Drosophila*. We show that this approach is robust to wide variations in video spatial and temporal resolution and that it can be used to measure leg segment motions during a variety of locomotor and grooming behaviors. FlyLimbTracker, the software implementation of this method, is open-source and our approach is generalizable. This opens up the possibility of tracking leg movements in other species by modifications of underlying active contour models.

**Author Summary:** In terrestrial animals, including humans, fundamental actions like locomotion and grooming emerge from the displacement of multiple limbs through space. Therefore, precise measurements of limb movements are critical for investigating and, ultimately, understanding the neural basis for behavior. The vinegar fly, *Drosophila melanogaster*, is an attractive animal model for uncovering general principles about limb control since its genome and nervous system are easy to manipulate. However, existing methods for measuring leg movements in freely behaving *Drosophila* have significant drawbacks: they require complicated experimental setups and provide limited information about each leg. Here we report a new method - and provide its open-source software implementation, FlyLimbTracker - for tracking the body and leg segments of freely behaving flies using only computational image processing approaches. We illustrate the power of this method by tracking fly limbs during five distinct walking and grooming behaviors and from videos across a wide range of spatial and temporal resolutions. Our approach is generalizable, allowing researchers to use and customize our software for limb tracking in *Drosophila* and in other species.

## Introduction

Many terrestrial animals rely on complex limb movements to locomote, groom, court, mate, and fight. Discovering how these and other fundamental behaviors are orchestrated by the nervous system will require manipulations of the genome and nervous system as well as quantitative measurements of behavior. The vinegar fly, *Drosophila melanogaster*, is an attractive model organism for uncovering the neural and genetic mechanisms underlying behavior. First, it boasts formidable genetic tools that allow experimenters to remotely activate, silence, visualize and modulate specific gene function in identified neurons [1]. Second, a number of sophisticated methods have been developed that permit robust tracking of *Drosophila* body movements – a promising set of tools for high-throughput screens [2-7].

By contrast, similarly robust methods with the precision required to semi-automatically track leg segments are largely absent. State-of-the-art approaches suffer from several drawbacks. For example, the most precise methods require the manual placement of visible markers on tethered animals [8] as well as sophisticated fluorescence-based optics (for another example in cockroaches see [9]). Marking insect leg segments is a time-consuming process that limits experimental throughput. On the other hand, the most high-throughput approach for marker-independent leg tracking in freely behaving *Drosophila* uses complex optics to measure Total-Internal-Reflection Fluorescence (TIRF) when the distal leg tips (claws) of walking animals scatter light transmitted through a transparent floor [10]. Although this method can resolve the claws of each leg it cannot detect their segments. Thus, it provides only binary information about whether or not a leg is touching the surface and cannot resolve the velocity of legs during swing phases, stance adjustments, or non-locomotive limb movements such as reaching [11] or grooming [12].

Here we describe a new method that permits semi-automated, marker-free tracking of the body and leg segments of freely walking *Drosophila.* We implement this method in an open source software plugin for Icy named FlyLimbTracker. Our approach uses active contours (i.e., snakes) to process objects in high-frame-rate image sequences. Thus, it does not require complicated optical setups. While there are a number of active contour algorithms [13], here we use parametric spline-snakes. These global-purpose, semi-automated image segmentation algorithms are typically used in two steps. First, the user roughly initializes a curve to a feature in an image (e.g., a fly’s body or leg). Second, the curve’s shape is automatically optimized to fit the boundaries of the object of interest. Therefore, segmentation algorithms using spline-snakes are composed of two major components: a *spline curve* or *model* that defines how the snake is represented in the image, and a *snake energy* that dictates how the curve is deformed in the image plane during optimization. Spline-snake models have a number of advantages to other approaches: they are (i) composed of only a few parameters, (ii) very flexible, (iii) amenable to easy manual edits, and (iv) formed from continuously defined curves that permit refined data analysis. Such models have therefore become widely used for image segmentation in medium-throughput biological applications [14,15]. Using this approach, we show that FlyLimbTracker can semi-automatically track freely walking or grooming *Drosophila melanogaster* in video data that spans a wide range of spatial and temporal resolutions. FlyLimbTracker is written as a plug-in for Icy, an open-source, community-maintained, and user-friendly image processing environment for biological applications [16-18]. This makes it amenable to customization for behavioral measurements in other species.

## Materials and Methods Drosophila

### behavior experiments

We performed experiments using adult female *Drosophila melanogaster* of the *Canton-S* strain at 2-4 days post-eclosion. Flies were raised on a 12 h light:12 h dark cycle at 25**°**C. Experiments were performed in the late afternoon Zeitgeber time after flies were starved for 4-6 h in humidified 25**°**C incubators.

During experiments, we placed flies in a custom designed acrylic arena (pill shaped: 30 mm x 5 mm x 1.2 mm) illuminated by a red ring light (FALCON Illumination MV, Offenau, Germany). We captured behavioral video using a high-speed (236 frames-per-second), high-resolution (2560 x 918 pixels) camera (Gloor Instruments, Uster Switzerland).

### Automated body and leg tracking

FlyLimbTracker is implemented in Java as a freely available plug-in for Icy, a cross-platform, multi-purpose image processing environment [16]. Briefly, FlyLimbTracker performs leg segment tracking in several steps. First, the user is asked to manually initialize the position of a fly’s body and leg segments in a single frame of the image sequence. This information is combined with image features to propagate body and leg segmentation to the frames immediately preceding, or following this first frame. At any time, the user can stop, edit, and restart automated segmentation. Manual corrections are taken into account when tracking is resumed.

To perform image segmentation, FlyLimbTracker uses active contour models (i.e., snakes). A snake [19] is defined as a curve that is optimized from an initial position - usually specified by the user - toward the boundary of an image object. Evolution of the curve’s shape results from solving an optimization problem in which a cost function, or snake energy, is minimized. Thus, snakes are an effective hybrid, semi-automated algorithm in which user interactions define an initial position from which automated segmentation proceeds [20,21]. Specifically, FlyLimbTracker first uses a *closed* snake to segment the *Drosophila* body into a head, thorax, and abdomen. Then, *open* snakes are used to model each of the fly’s legs. Manual mapping of these snakes onto the fly in an initial frame is the basis for subsequent tracking.

### Drosophila *body model*

We designed a custom snake model to segment and track the *Drosophila* body. In our model, the fly’s body is defined as a 2-dimensional closed curve **r**:

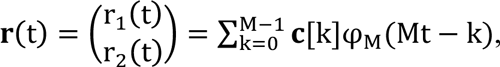

with *t* ∈ [0, *M*), where **c**[*k*] = {(c_1_[*k*]c_2_[*k*])^T^}_*k*∈ℤ_ is an *M*-periodic sequence of control points and 
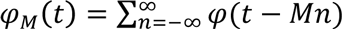
 the *M*-periodization of a basis function *φ*. For a thorough description of the spline snake formalism, see [13]. The proposed model for the body of the fly consists of an *M*=18 nodes snake using the ellipse-reproducing basis [22]

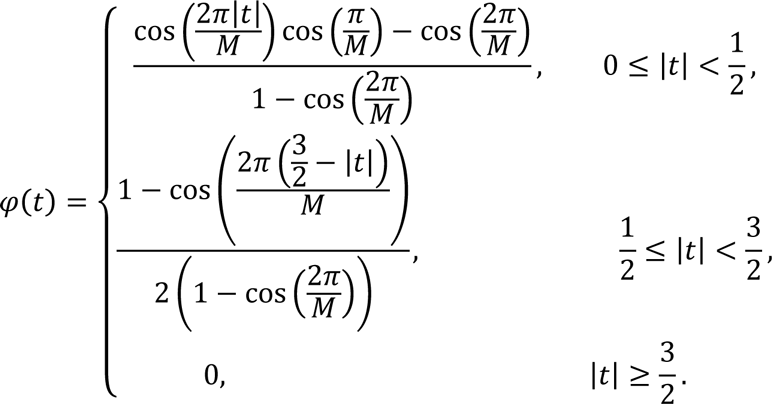

To optimize the snake automatically from a coarse initialposition to the precise boundaries of the fly’s body, we define a snake energy composed of three elements:

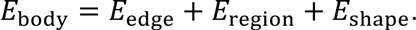

The first element *E*_edge_ is an edge-based energy term relying on gradient information to detect the body contour, which is formally expressed as

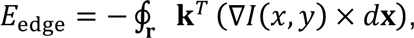

where *d***x** is the infinitesimal vector tangent to the snake, ∇*I*(*x*, *y*) the in-plane gradient of the image at position (*x*, *y*), and **k** = (0, 0, 1) is the vector orthonormal to the image plane. The energy term is negative since it has to be minimized during the optimization process. Using Green’s theorem, we can transform the line integral into a surface integral:

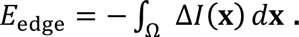

The second term, *E*_region_, is a region energy term that uses region statistics to segment the object from the background. Specifically, it is computed as the intensity difference between the region enclosed by the snake and the region surrounding it, as

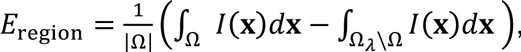

where *I* is the image and |Ω| the signed area of the snake, which is defined as

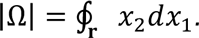

Minimizing this term encourages the snake to maximize the contrast between the area it encloses and the background. For more details about the edge and region energy derivations, see [23,24].

Finally, the last term, *E*_shape_, corresponds to the shape-prior energy contribution detailed in [25]. This term measures the similarity between the snake and its projection on a given reference curve. It therefore encourages the convergence of the contour to an affine transformation of the reference shape. The smoothness and regularity of the reference are preserved. Moreover, this term prevents the formation of loops and aggregation of nodes during the optimization process. In our case, the reference shape is a symmetric 18-node fly body contour (Fig. 1A,F).

**Figure 1.**
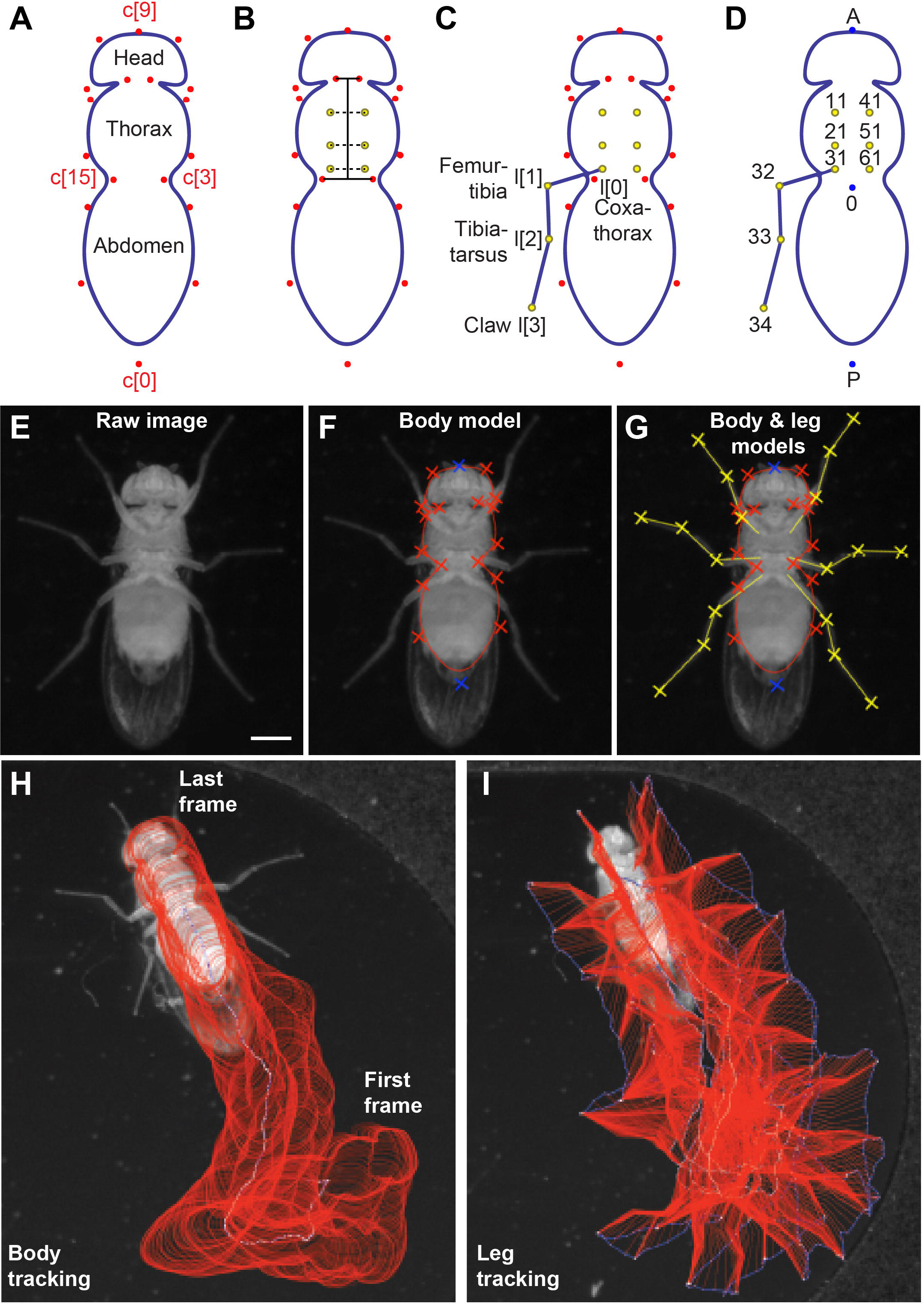
FlyLimbTracker uses active contour models to annotate the *Drosophila* body and legs. **(A)** The body model is a closed snake consisting of 18 control points (**c**[0] to **c**[17]). Control points **c**[0] and **c**[9] correspond, respectively, to the posterior-most position on the abdomen and the anterior-most position on the head. All other control points are symmetric along the anteroposterior axis of the body (e.g., control points **c**[3] and **c**[15]). **(B)** Six leg anchor positions (yellow) between the coxa and thorax are defined empirically based on a linear combination of distances from the head-thorax boundary, the thorax-abdomen boundary, and a distance from the thoracic midline. These positions are then shifted depending on how the body model is optimally deformed to fit the contours of a specific animal. **(C)** The leg model consists of four control points including a thorax-coxa attachment **I**[0], the femur-tibia joint **I**[1], the tibia-tarsus joint **I**[*2*], and the pretarsus/claw **I**[3]. For simplicity, control points for only a single leg are shown. **(D)** In sum, 27 positions are calculated for each fly per frame: a centroid (0), anterior point (A), posterior point (P), as well as the body anchor, first intermediate, second intermediate and tip for each of the six legs. Our data labeling convention is as follows. Right and left legs are numbered 1 to 3 (front to rear) and 4 to 6 (front to rear), respectively. Each leg has four control points labeled 1 to 4 in the units digit that correspond the body anchor (1), leg joints (2 and 3), and claw (4). In each label, the leg number is shown in the tenths digit and the control point in the units digit. For example, the label “11” refers to the body anchor of the right prothoracic leg 1. For simplicity, only the control points for leg 3 are shown. **(E)** An example raw image of the ventral surface of a fly used for segmentation. **(F)** This image is first segmented using the parametric body snake consisting of 18 control points (red and blue crosses). **(G)** Subsequently, leg segmentation is initialized through automatic tracing from body anchor points to user-defined leg tips. From this initialization, an annotation is performed using open snakes consisting of four control points (yellow crosses). (**H**) Body and (**I**) leg segment tracking annotation for flies during a 455-frame (1.93 s) sequence. Annotation results (red) and the centroid in **H** or leg tip positions in **I** (blue) for each frame are overlaid.

To automatically optimize the snake, we modified the position of the control points by minimizing the energy using a Powell-like line-search method [26], a standard unconstrained optimization algorithm that converges quadratically to an optimal solution. First, one direction is chosen depending on the partial derivatives of the energy, which is computed using finite differences. Second, a one-dimensional minimization of the energy function is performed in the selected direction. Finally, a new direction is chosen using the partial derivatives and enforcing conjugation properties. These steps are repeated until convergence. The final configuration of the control points provides an accurate description of the orientation and size of the fly body.

In practice, the algorithm depends on initial user input to coarsely locate the fly in a frame of the image sequence. Following a single mouse click, a two-step multiscale optimization scheme inspired by [24] is initiated. A spherical active contour composed of 3-control points is first created, centered at the mouse position. This snake is optimized using *E*_edge_ + *E*_region_ to form an elliptic curve surrounding the fly. In this way, the major axis of the elliptical snake will be aligned with the anteroposterior axis of the fly, and the minor axis will be perpendicular to it.

The 3-point elliptical snake fit to the body of the fly can be expressed as follows [23]:

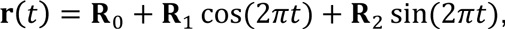

where

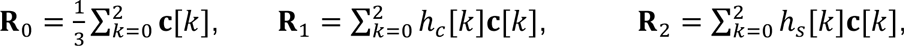

and

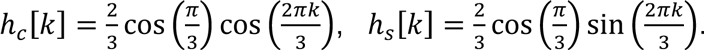

Relating this to the general parametric equation of an ellipse of major axis *a*, minor axis *b*, and center (*x*_c_ *y*_c_)^T^ allows us to extract the parameters of the 3-control point snake fit to the fly’s body. Namely, (*x*_c_ *y*_c_)^T^ = **R**_0_, a = max (‖**R**_1_‖, ‖**R**_2_‖) and b = min (‖**R**_1_‖, ‖**R**_2_‖). By knowing *a*, the orientation of the ellipse in the image can be computed.

The ellipse fit is then replaced by an 18-node fly-shaped closed snake that has been rotated and dilated to match the ellipse’s length and orientation (Fig. 1A). An ambiguity results since two potential snake models can be initialized for a given ellipse, with opposite anteroposterior axis orientation. To resolve this ambiguity, both potential snake orientations are optimized on the image using *E*_body_ in addition to *E*_edge_ and *E*_region_. The solution with the lowest cost (i.e., energy value at convergence) is used.

### Drosophila *leg model*

Once the fly’s body is properly segmented, open snake models for each of its legs are then added. First, the positions of leg coxa-thorax attachment points (hereafter referred to as *anchors*) are automatically computed based on the body segmentation. The location of the six leg anchors with respect to the reference body model have been empirically determined as linear combinations of three axes defined by the head-thorax junction, the thorax-abdomen junction and the thorax length (Fig. 1B). These locations are then adapted according to an individual fly-specific deformation of the body model.

User input is required to initialize the positions of each leg prior to tracking. Initialization is based on a single click for each leg: the user indicates the claw (hereafter referred to as *tip*) of each leg through mouse-clicks on the selected frame. The click location is assigned to the most likely body anchor using a probabilistic formulation based on the distance and intersection with the fly’s body model and that of other leg models. Once a leg tip and a leg anchor have been paired, a dynamic programming method [27] is initiated to automatically trace the leg from the anchor to the tip. To facilitate this process, the fly’s legs are enhanced by processing the segmented image frame using a ridge detector [28].

Dynamic programming is a method that yields the globally optimal solution for a given separable problem. In particular, it can be used to implement algorithms solving shortest path problems. Dynamic programming relies on a graph-based representation: the shortest path is represented as a sequence of successive nodes in a graph that minimize a cost function. To trace a leg from its anchor to its tip, we build a graph by interpolating image pixels along the two axes using a straight segment linking the anchor to the tip (axis **k**) and its normal vector (axis **u**). The cost of the path at index *k* + 1 along axis **k** is then given by:

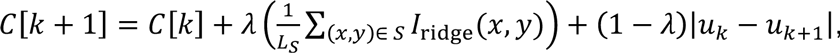

where *C*[*i*] is the cost of the path at location *i* on axis **k**, *S* is the collection of image pixels (*x*, *y*) in the segment between node (*k*, *u_k_*) and (*k* + 1, *u*_*k* + 1_), *L_s_* is the pixel length of this segment, *I*_ridge_ is the ridge-filtered version of current frame, and *λ* ∈ [0, 1] is a weighting coefficient. The first term corresponds to a discretized integral of the image in the segment linking nodes *k* and *k* + 1, and therefore tends to favor paths going through low pixel values. The second term is composed of the distance along axis **u** between two successive nodes. As a result, the optimal path follows relatively bright (or dark) regions in the image with respect to the background, while retaining a certain level of smoothness. The relative contribution of each term is determined by *λ*.

In contrast to body segmentation, leg segmentation uses open rather than closed snakes. Fly legs are parameterized by a curve composed of *M* = 4 control points (Fig. 1C,G). For each leg, the body anchor, **l**[0], is considered fixed. The discrete path obtained through dynamic programming is used to initialize the leg snake. The rationale behind this two-step procedure is two-fold. First, dynamic programming is very robust and can therefore effectively trace the leg from a body anchor to its tip. However, since it is a discrete approach, it is computationally expensive. By contrast, snake-based methods are more likely to diverge when initialized far from their target but are computationally inexpensive since only a few control points need to be stored to characterize a given curve. Therefore, we combined these approaches by first finding a path to define each leg using dynamic programming and then transforming this path into a parametric curve for optimization. The parametric representation of the leg snake curve is defined as

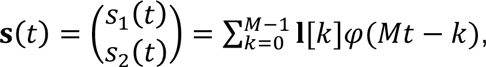

where *t* ∈ [0, *M* − 1] and **I**[*k*] = {*l*_1_[k]*l*_1_[k])^T^}_*k*∈ℤ_ are the leg snake control points. Since *Drosophila* legs are composed of relatively straight segments between each joint, we use linear splines as basis functions *φ*(*t*). The leg control points are therefore linked through linear interpolation and each control point has a unique identifier that can be used for subsequent data processing (Fig. 1D). Figure 1E-G illustrates the full process of taking a single raw image (Fig. 1E) and using active contours to segment the body (Fig. 1F) and legs (Fig. 1G).

### Segmentation propagation (tracking)

High frame-rate videos ensure that the displacement of a fly’s body between successive frames is small. FlyLimbTracker takes advantage of this fact to propagate body and leg snakes from one frame to the next during tracking. The body snake in frame *t+1* is therefore segmented by optimizing a contour initialized as the corresponding snake from frame *t* using the body snake energy previously described. This approach is sufficient to obtain good segmentation provided that there is some overlap between the animal’s body in frames *t* and *t+1*.

Compared with the body, leg displacement can be larger between frames. Therefore, leg snakes require a more sophisticated algorithm to be propagated during tracking. First, the anchor of each leg is automatically computed from the newly propagated fly body. Since each leg is modeled as a 4-node snake, the three remaining leg snake control points are optimized using the snake energy

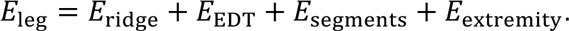

The first term corresponds to the integral along the leg in the current frame filtered by a ridge detector [28], i.e.,

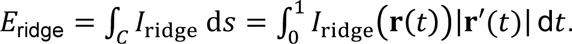

Analogous with the first term, the second term is computed as the integral along the leg of the Euclidean distance transform (EDT, [29]) in the current frame where

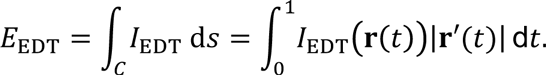

Each of the linear segments comprising a fly’s legs should be roughly constant in length across a video, aside from changes introduced by projecting the three-dimensional legs onto two-dimensional images. Taking this consistency into account, the third term of the leg energy penalizes solutions for which the leg joint positions result in leg segments whose lengths vary considerably from one frame to the next. This prevents unrealistic configurations of the leg joints that yield excessively long leg segments compared with neighboring annotated frames.

Finally, the fourth term is used to determine the leg tip position at time *t*, denoted **I**_t_[3]. Since the distal tip of the leg may move considerably between successive frames, we designed a dedicated energy term to attract the tip toward candidate locations in the image. These candidate locations are defined by minima after the image is filtered using a Laplacian-of-Gaussian (LoG, [30]). A potential map of tip candidates is then created according to:

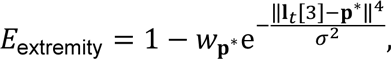

where

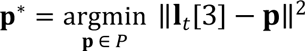

is the tip candidate closest to **I**_*t*_[3] 
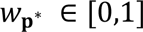
 its associated weight, and *σ*^2^ a fixed parameter determining the width of the attraction potential of the tip candidates. The weight 
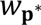
 is a measure of how tip-like **p**^*^ is, and is computed based on the magnitude of the LoG filter response. A strong weight results in a deeper potential, and is therefore more likely to attract **I**_*t*_[3].

In summary, the four anchor points characterizing each leg are propagated as follows. First, the leg body anchors are determined using the body model. Second, the remaining three control points (two leg joints and tip) are shifted by optimizing a cost function that incorporates both image information (*E*_ridge_, and *E*_EDT_) and a smoothness constraint (*E*_segmetns_). Finally, the tip is further constrained using an estimation of how tip-like the image is at candidate locations.

### Data output

Once the full image sequence is annotated, data can be extracted as a CSV file for each fly. These measurements include the locations of three reference points on the fly’s body (A, P, and 0), as well as each of the legs’ anchor points (see Fig. 1D for the labeling convention).

FlyLimbTracker is linked to Icy’s Track Manager plugin (Publication Id: ICY-N9W5B7) via the *extract tracks* buttons (see interface description in the Appendix), allowing additional data to be extracted. In particular, segmentations of the fly’s body (Fig. 1H) and legs (Fig. 1I) can be visualized across the entire sequence, illustrating their entire trajectories. Each individual control point of the leg snakes or the body snake’s centroid can be independently visualized. Note that tracks are also numbered according to the labeling convention in Fig. 1D.

### Software and data availability

User instructions, FlyLimbTracker software, and sample data can be found at: http://bigwww.epfl.ch/algorithms/FlyLimbTracker/

## Results

FlyLimbTracker performs semi-automated body and leg tracking. First, the user manually initializes the positions of the fly’s body and leg segments in a single, arbitrarily chosen frame of the image sequence (Fig. 2A). These manual annotations are then used to automatically propagate segmentation to prior, or subsequent frames (Fig. 2B). During automated segmentation, the user can interrupt tracking to correct errors (Fig. 2C). When FlyLimbTracker is restarted, the automated segmentation continues, taking into account these user edits.

**Figure 2.**
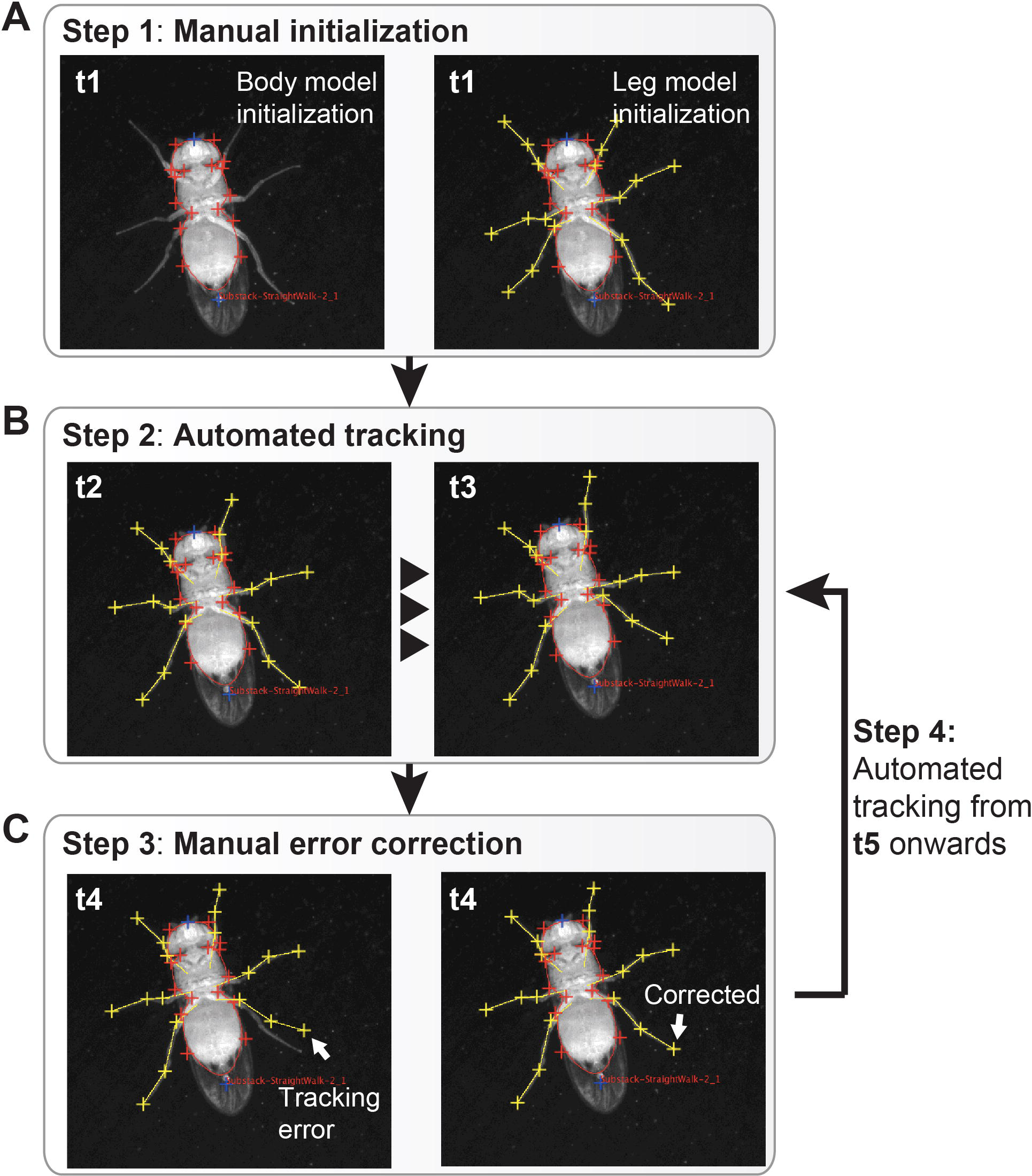
FlyLimbTracker workflow. **(A)** The user manually indicates the approximate location of the fly’s body in an arbitrarily chosen video frame (t1). FlyLimbTracker then optimizes a closed active contour model that encapsulates the fly’s body in the correct orientation. The user then manually indicates the location of each leg’s tip. FlyLimbTracker then optimizes an open active contour model that runs across the entirety of each leg. **(B)** The user then runs FlyLimbTracker’s automatic tracking algorithm to propagate body and leg models to subsequent video frames (or prior frames if run in reverse). **(C)** Either during or after automated tracking, the user can look for tracking errors. After manually correcting these errors, the user can re-run automatic tracking. In each image, the frame number is indicated.

### Algorithm robustness

FlyLimbTracker can be used to segment and track fly bodies and legs in videos spanning a wide range of spatial and temporal resolutions. Resolution determines the nature of the annotation process: high-resolution data tracking is more automated, while low resolution data requires more user intervention. To quantify the dependence of computing time and the number of user interventions on data quality, we systematically varied the spatial and temporal resolutions of videos featuring five common *Drosophila* behaviors: walking straight, turning, foreleg grooming, head grooming, and abdominal grooming. Raw videos were originally captured at 236 fps and at 2560 x 918 pixel resolution (Supplementary Videos 1-5).

First, we studied FlyLimbTracker’s robustness to variations in spatial resolution. We down-sampled each of the five videos by a factor of *N*, where *N* × *N* pixels were averaged. This resulted in image sequences *N* times smaller along both spatial dimensions but with an identical temporal resolution of 236 fps (Fig. 3A). Alternatively, to vary temporal resolution, we down-sampled each video by a factor of *N*, where only one frame from every *N* was retained. This resulted in image sequences of varying temporal resolution but consistently high spatial resolution of 2560 x 918 pixels (Fig. 3B).

**Figure 3.**
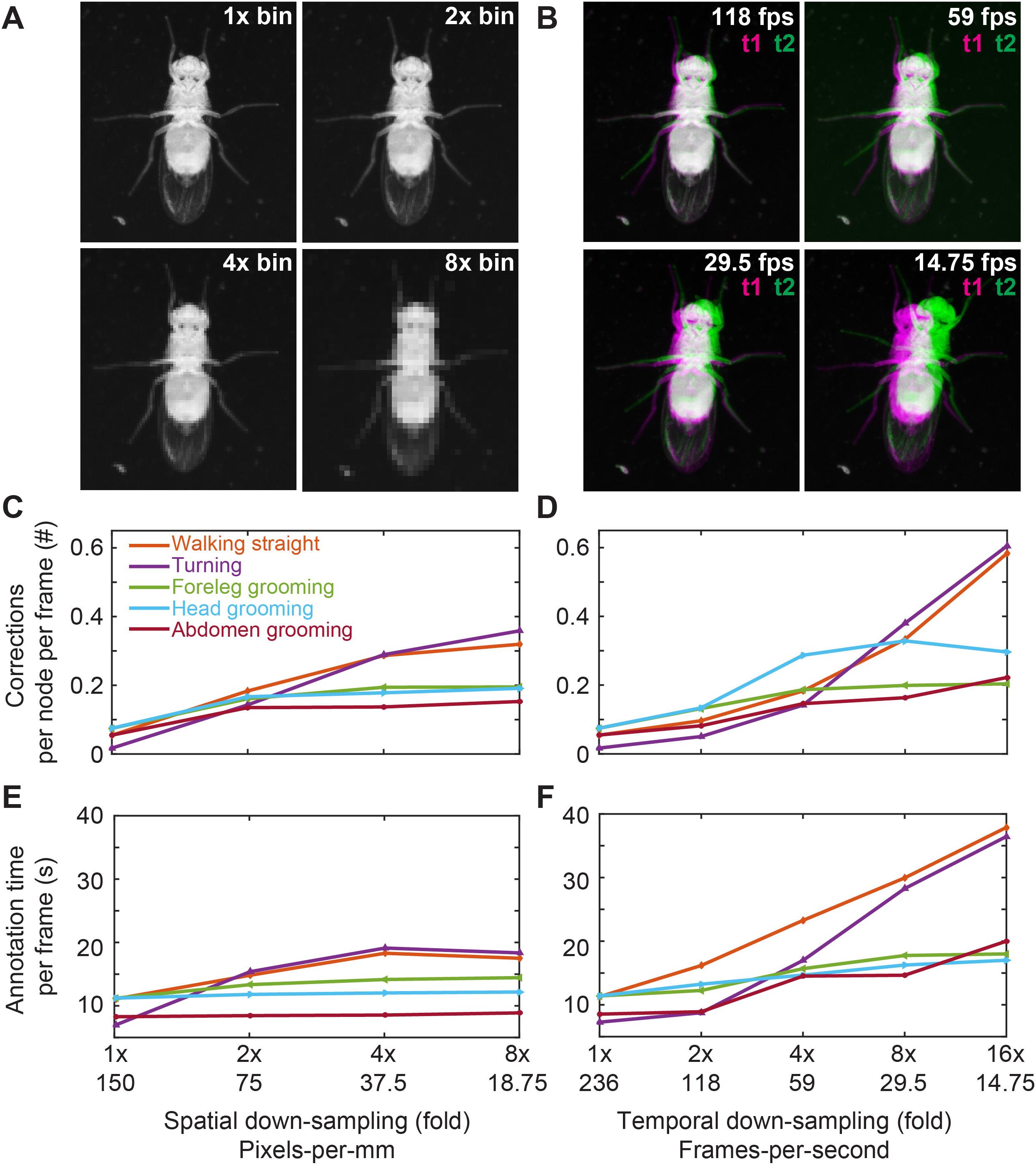
Sensitivity of leg tracking to changes in spatial or temporal video resolution. **(A)** Sample video image (top-left) after 2x (top-right), 4x (bottom-left), or 8x (bottom-right) spatial down-sampling. **(B)** Representations of the difference between successive images (t1 and t2 overlaid in magenta and green, respectively) for different frame rate videos after temporal down-sampling. **(C-D)** The number of corrections required per node per frame as a function of spatial resolution **(C)**, or temporal resolution **(D)**. **(E-F)** The average time required to annotate a single frame as a function of spatial resolution **(E)**, or temporal resolution **(F)**. In **C-F**, data for videos depicting a fly walking straight, turning, grooming its forelegs, head, or abdomen are shown in orange, purple, green, cyan, and red, respectively.

For each movie, body and leg snakes were manually initialized using the first image frame. Segmentation was then automatically propagated forward through the remainder of the image sequence. Whenever the automated tracker made a mistake, the process was interrupted and the user manually corrected the error. Automated tracking was then restarted from this frame until the next mistake was observed. In all cases, automated body tracking did not require manual intervention. Therefore, we only took note of manual corrections in leg snake annotation.

To quantify FlyLimbTracker’s performance across this range of spatial and temporal resolutions, we calculated two normalized quantities. First, we calculated the average number of manual corrections per node per frame (Fig. 3C-D). To do this, we measured the total number of user interventions while processing an image sequence and normalized this quantity by *T*×6×3, where *T* is the number of frames, each of which contains eighteen free parameters: six legs with three editable control points each. As a second metric we quantified the average time required to annotate a single image frame (Fig. 3E-F). To do this, we recorded the total time required to annotate an image sequence and divided this value by the total number of frames. This normalized quantity combines both the computing time required for automated annotation as well as the time required to manually correct annotation errors.

Overall, we observed that reducing spatial (Fig. 3A,C,E), or temporal (Fig. 3B,D,F) resolution resulted in an increase in the number of manual interventions (Fig. 3C-D) as well as a longer time required for annotation (Fig. 3E-F). While the numbers of corrections were similar for equivalent amounts of down-sampling (up to 8-fold), annotation time was appreciably longer for straight walking and turning. This reflects the importance of having overlapping images in successive frames for automated tracking: a feature that may be less common during locomotion where the position of a leg can vary substantially within a walking cycle. Notably, in a number of other cases (e.g., grooming), the annotation time per frame flattens across spatial and temporal resolutions. This is probably due to the trade-off between automated processing and manual correction times. Resolution strongly influences the computing time required for automated tracking: smaller or fewer images can be processed more quickly. However, as resolution decreases, user interventions required to correct errors begin to dominate annotation time required to annotate each frame.

### Visualization and analysis of leg segment tracking data

FlyLimbTracker provides a user-friendly interface that allows body and leg segment tracking data to be exported in a CSV file format, simplifying data analysis and visualization. We illustrate three representations of body and leg tracking data for annotated videos of the five behaviors previously described (Supplementary Videos 6-10). First, within FlyLimbTracker itself, leg joint and/or body trajectories can be displayed overlaid upon the final raw video frame (Fig. 4A_1_-E_1_). This representation provides a way to project time-varying data onto a static image and illustrates the symmetric or asymmetric limb motions that control straight walking/grooming or turning, respectively. Second, leg segment trajectory data can be exported and processed externally (e.g., using Matlab or Python). These data can be rotated along with the fly’s frame of reference (Fig. 4A_2_-E_2_) for a direct comparison of leg segment movements between distinct actions. A similar approach has been used to visualize how neurogenetic perturbations influence claw movements during locomotion [10], but can now be used to study the effects of these manipulations on other previously inaccessible leg segments and behaviors (e.g., grooming or reaching). In a third visualization, the speeds of each claw can be plotted to provide an exceptionally detailed characterization of locomotor gaits (Fig. 4A_3_-B_3_), or grooming movements in stationary animals (Fig. 4C_3_-E_3_).

**Figure 4.**
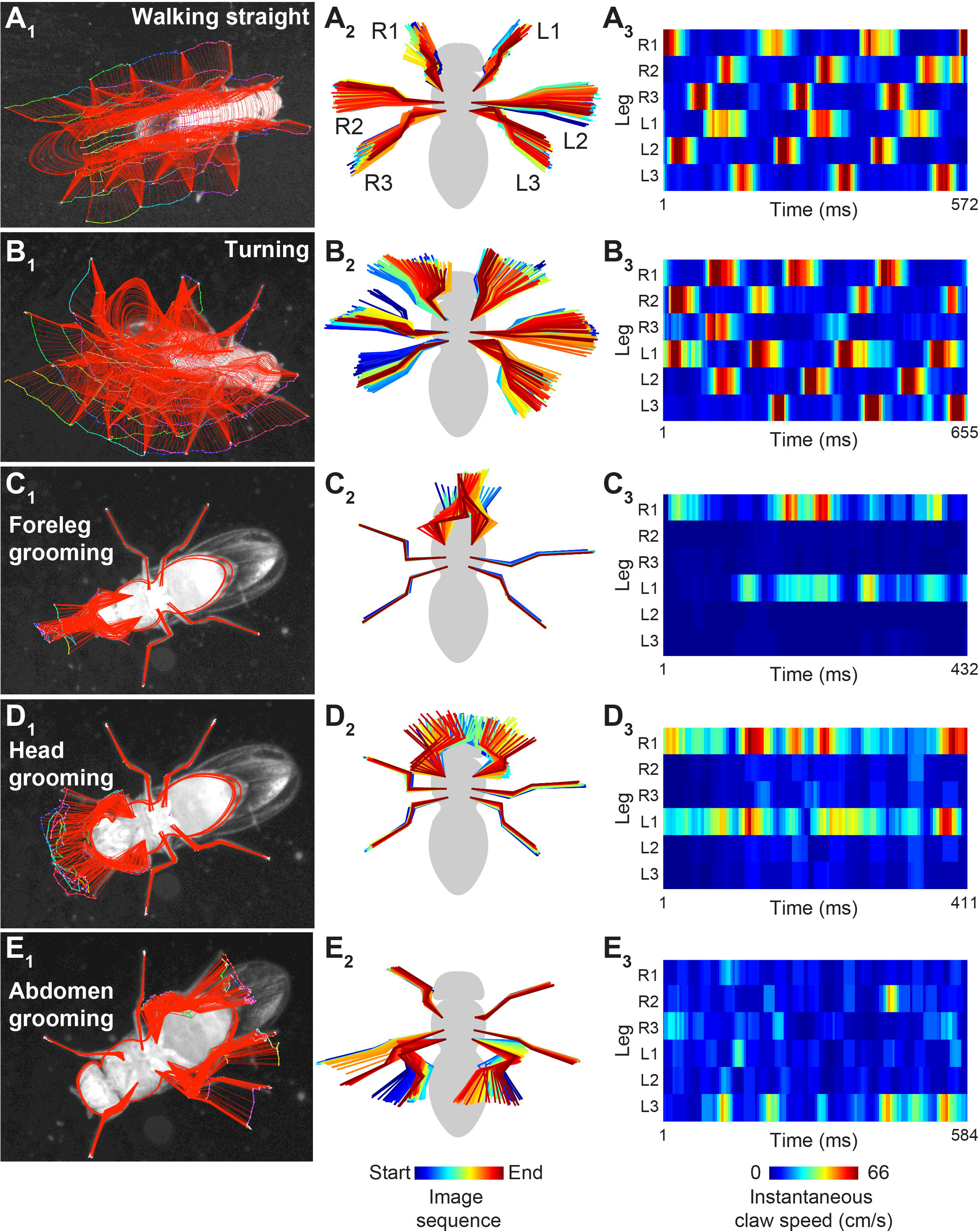
Analysis and visualization of FlyLimbTracker leg tracking data. Visualizations of leg segment annotation results for videos of a fly **(A)** walking straight, **(B)** turning, **(C)** grooming its forelegs, (**D**) grooming its head, or (**E**) grooming its abdomen. **(A_1_-E_1_)** Leg segmentation results (red) and joint positions (color-coded by frame number) are overlaid on the final frame of the image sequence. **(A_2_-E_2_)** Leg segment trajectories are rotated and color-coded by frame number. This permits alignment and comparison of leg movements across different datasets. **(A_3_-E_3_)** The instantaneous speeds of each leg tip (claw) are color-coded.

## Discussion

Existing methods for tracking insect leg segments rely on sophisticated optical equipment and/or laboriously-applied leg markers, often in tethered animals [8-10]. While these approaches are extremely valuable, they may potentially disrupt natural behaviors and cannot report the motions of multiple joints in untethered animals. Here we have introduced a method that uses computer-vision techniques to address these technical barriers. The software implementation of this approach, FlyLimbTracker, permits semi-automated tracking of body and leg segments in freely behaving *Drosophila*. Use of FlyLimbTracker only requires a single high-resolution, high-speed camera and does not require prior marking of leg segments. Additionally, it can be used with video data across a range of spatial and temporal resolutions, permitting a flexible blend of automated and manual annotation. Importantly, when automation has difficulty segmenting low quality data, FlyLimbTracker remains a powerful tool for manual leg tracking annotation since it uses easily manipulated spline-snakes and provides an interface for user-friendly data import and export.

The open-source nature of FlyLimbTracker can facilitate community-driven improvement and customization of the algorithm. We can envision a number of improvements moving forward. First, tracking currently requires overlap of a fly’s body between successive frames. This constraint places a lower bound on video temporal resolution and could be improved by using, for example, nearest-neighbor matching approaches like the Hungarian algorithm [31] to link segmentation control points between successive frames. Second, additional leg control points may be added to FlyLimbTracker to more precisely annotate thorax-coxa-trochanter segments. Third, FlyLimbTracker’s requirement of user initialization, makes it only semi-automated and restricts batch processing of multiple videos for high-throughput data analysis. This may be overcome using additional prior information to automatically identify and optimize body snakes. Fourth, FlyLimbTracker’s snake-based approach to tracking could easily be adapted for the study of other species (e.g., mice, stick insects, and cockroaches) by modifying the shape of snake models.

## Acknowledgements

We thank Cédric Vonesch, Michael Rusterholz, and Loic Perruchoud for early contributions to the body tracking algorithm.

## Supporting Information

Supporting Video Legends

Raw videos used for sensitivity analyses (Fig. 3) and visualization (Fig. 4):

Video 1 – A fly walking straight.

Video 2 – A fly turning.

Video 3 – A fly grooming its forelegs.

Video 4 – A fly grooming its head.

Video 5 – A fly grooming its abdomen.

Video 6 – A fly walking straight (video 1), annotated using FlyLimbTracker.

Video 7 – A fly turning (video 2), annotated using FlyLimbTracker.

Video 8 – A fly grooming its forelegs (video 3), annotated using

FlyLimbTracker.

Video 9 – A fly grooming its head (video 4), annotated using FlyLimbTracker.

Video 10 – A fly grooming its abdomen (video 5), annotated using

FlyLimbTracker.

## Appendix

### User interface

FlyLimbTracker’s interface can be used in either basic or advanced mode. In the basic mode, only the name of the active image is visible. All parameters are hidden and only default parameter values are used. When switching to the advanced mode, all parameters become visible and can be adjusted by the user. Parameters that can be adjusted in the interface include:

- Image parameters

- <u>Channel</u>: for multichannel images (e.g., bright-field and fluorescence), this parameter selects the channel upon which segmentation is performed. In most cases, the bright-field channel should be selected.
- <u>Smoothing</u>: adjusts the width (standard deviation, in pixels) of a smoothing filter used to preprocess the image sequence. Larger values yield smoother images, but likely obscure details such as the fly’s legs. We recommend choosing a value approximately equal to the average width (in pixels) of the fly legs.
- <u>Subtract background</u>: performs background subtraction on the image sequence. The background model used is the median of each pixel across the whole image sequence. In practice, background subtraction is not desirable in datasets with a low signal-to-noise ratio since a fly’s legs typically have low contrast and can be smoothed out by median filtering.
- Body model parameters

- <u>Annotation method</u>: switches between automated and manual annotation of the body snake. Automated annotation is obtained by automatically optimizing the body snake from its initial, manually chosen position. Manual annotation relies exclusively on user interactions.
- <u>Energy trade-off</u>: adapts the relative importance of data fidelity (image-based) and regularization (shape-based) terms in the body snake energy. A fully image-based snake would be optimized using image information only, while a fully shape-based snake would be optimized to retain a fly’s shape regardless of the underlying image data. For data with low image quality the regularization term (shape-based) becomes more important.
- <u>Max iterations/immortal</u>: tunes the maximum number of iterations used to optimize the body snake. If *immortal* is chosen, the snake keeps evolving until it achieves convergence. Allowing the snake to be immortal usually yields better segmentation results, but significantly increases computing time. Conversely, a smaller number of iterations can estimate segmentation quickly, but not necessarily as effectively. Usually, 4000-5000 iterations provide a good trade-off between computing time and segmentation quality. However, this value should be customized according to data quality.
- <u>Freeze snake body</u>: when ticked, locks the control points of the fly body snake, which then appear as blue instead of red. In this setting, individual points cannot be further edited. This feature is useful when the fly body is properly initialized and edits are done on the legs only, as it prevents displacing body control points when trying to select a leg control point. However, it remains possible to translate, move or rotate the entire fly body.
- Leg model parameters

- <u>Annotation method</u>: switches between automated and manual segmentation of the fly’s legs. Although body segmentation and tracking is robust even for low resolution or low signal-to-noise ratio data, leg tracking is much more sensitive. Therefore, the user is given the option to restrict automation to body tracking. In the manual segmentation setting, the legs are simply propagated by translation along with body motion and can be manually adjusted post-hoc for each frame. This allows FlyLimbTracker to be a useful tool for annotating either low-quality or high-quality data.
- <u>DP trade-off</u>: determines the relative importance of data fidelity (bright) and regularization (straight) terms when performing dynamic programming (DP) to initialize the leg snakes. The algorithm tries to find the optimal path between a given leg anchor and tip by optimizing the trade-off between image intensity (bright) and straightness (straight). Relying on image brightness alone typically yields irregular movements of the fly’s legs since the algorithm becomes very sensitive to image noise (e.g., isolated pixels of high intensity). Conversely, relying on straightness alone yields, in the most extreme case, a straight line between the anchor and tip. Note that this parameter is only used when initializing a leg. It does not influence tracking.
- <u>Energy trade-off</u>: determines the relative importance of data fidelity (image-based) and regularization (sequence-based) terms for the leg snakes. A purely image-based leg snake is optimized using the image data only. This typically yields suboptimal solutions that are sensitive to image noise. Conversely, a fully sequence-based leg snake maximizes its resemblance to the corresponding leg snake from previously annotated frames and ignores image data. More importance should be given to sequence-based energy for low quality data when leg snake annotations are readily available.
- <u>Tip propagation mode</u>: determines the relative importance of data fidelity (image-based) and regularization (sequence-based) terms while tracking leg tips. We identify potential tips by searching for candidate locations in a neighborhood encompassing leg motions from previously annotated, neighboring frames. The final tip position is chosen as a trade-off between the position predicted by leg motion from previous annotated frames (sequence-based), and tip candidates identified by processing the current frame (image-based).
- <u>Max iterations/immortal</u>: tunes the maximum number of iterations used to optimize the leg snakes in a manner similar to how the same parameter is used to optimize the body snake.

In both basic and advanced modes, the upper part of the interface contains several menu items (Analyze, Save/Load and Help):

- <u>Analyze</u>: extracts measurements from the current body segmentation using Icy’s ROI Statistics plugin (Publication Id: ICY-W5T6J4).
- <u>Save/Load</u>: allows the user to export and save annotations to a CSV file format (see Output section below). This can also be used to reload previously saved CSV annotations.
- <u>Help</u>: contains information about the plugin version (About), and a link to FlyLimbTracker’s online documentation page (Documentation (online)).

Finally, several action buttons are located on the lower part of the interface. These are split into three sections.

- <u>Fly shape editing</u>: the left button enables movement of individual control points. The middle and right buttons, respectively, enable resizing and rotation of the body and leg snakes.
- <u>Snake action</u>: automatically optimizes the snake at its current position (left button), or deletes it (right button). Note that both actions are applied to the body snake and all leg snakes simultaneously. If annotation methods for body or leg snakes are set to *manual*, the corresponding snakes are left unmodified.
- <u>Tracker action</u>: performs backward (left button) or forward (center-left button) tracking, interrupts tracking (center-right button), or extracts/displays tracks (right button) using Icy’s Track Manager plugin (Publication Id: ICY-N9W5B7). The tracking algorithm is implemented to allow backward and forward tracking, giving the user flexibility to initialize tracking at any frame of the image sequence. If any of the body or leg snakes are set to manual annotation, the forward and backward tracking buttons will only propagate current annotations to the next or previous frame, respectively. If all snakes are set to automated annotation, tracking will be performed in the selected direction until the end/beginning of the image sequence is reached, unless it is manually halted using the tracking interruption button.

